# Female fibroblast activation is estrogen-mediated in sex-specific 3D-bioprinted pulmonary artery adventitia models

**DOI:** 10.1101/2025.01.17.633670

**Authors:** Mikala C. Mueller, Rachel Blomberg, Alicia E. Tanneberger, Duncan Davis-Hall, Keith B. Neeves, Chelsea M. Magin

## Abstract

Pulmonary arterial hypertension (PAH) impacts male and female patients in different ways. Female patients exhibit a greater susceptibility to disease (4:1 female-to-male ratio) but live longer after diagnosis than male patients. This complex sexual dimorphism is known as the estrogen paradox. Prior studies suggest that estrogen signaling may be pathologic in the pulmonary vasculature and protective in the heart, yet the mechanisms underlying these sex-differences in PAH remain unclear. PAH is a form of a pulmonary vascular disease that results in scarring of the small blood vessels, leading to impaired blood flow and increased blood pressure. Over time, this increase in blood pressure causes damage to the heart. Many previous studies of PAH relied on male cells or cells of undisclosed origin for *in vitro* modeling. Here we present a dynamic, 3D-bioprinted model that incorporates cells and circulating sex hormones from female patients to specifically study how female patients respond to changes in microenvironmental stiffness and sex hormone signaling. Poly(ethylene glycol)-alpha methacrylate (PEGαMA)-based hydrogels containing female human pulmonary artery adventitia fibroblasts (hPAAFs) from idiopathic PAH (IPAH) or control donors were 3D bioprinted to mimic pulmonary artery adventitia. These biomaterials were initially soft, like healthy blood vessels, and then stiffened using light to mimic vessel scarring in PAH. These 3D-bioprinted models showed that stiffening the microenvironment around female IPAH hPAAFs led to hPAAF activation. On both the protein and gene-expression levels, cellular activation markers significantly increased in stiffened samples and were highest in IPAH patient-derived cells. Treatment with a selective estrogen receptor modulator reduced expression hPAAF activation markers, demonstrating that hPAAF activation is a one pathologic response mediated by estrogen signaling in the vasculature, validating that drugs currently in clinical trials could be evaluated in sex-specific 3D-bioprinted pulmonary artery adventitia models.

## 1. Introduction

Pulmonary arterial hypertension (PAH) is a chronic, progressive fibrotic pulmonary vascular disease characterized by overactive pulmonary arterial tissue remodeling, which causes stiffening and narrowing of the pulmonary vasculature. This remodeling increases vascular resistance, causing the heart to work harder to pump blood through the lungs, and eventually leads to right ventricular heart failure.^1^ While some forms of PAH can be linked to genetic factors, idiopathic PAH (IPAH) is far more common, accounting for 30-48% of PAH cases.^2^ There is higher incidence of IPAH in females patients; however, females tend to live longer than their male counterparts after diagnosis. The underlying causes of these sex-differences remain largely unknown.^3–6^ Pulmonary artery adventitial fibroblasts (PAAFs) play a key role in early IPAH pathogenesis by depositing excess extracellular matrix (ECM) during sustained activation responses that contribute to tissue stiffening and disease progression, but how these critical cells contribute to sex differences in IPAH is critically understudied.^7–9^ Engineering sex-specific models of human PAAF (hPAAF) activation could create powerful tools for advancing our understanding of and treatments for IPAH.^10, 11^

Females are four times more likely to develop IPAH than males and are diagnosed with IPAH younger (median age of development of 36 ± 15 years).^1, 3–5, 12^ Despite females being more likely to develop IPAH, males tend to have worse ventricular function and survival rates. Studies have shown that increased estrogen levels promote IPAH development by causing pulmonary vasculature injury, but estrogen can also protect against severe disease progression by increasing right ventricle function in IPAH.^3–5, 13^ This phenomenon is referred to as the estrogen paradox. Different metabolites of estradiol have been found to have contradictory effects on the pulmonary arteries, where 16α-hydroxyoestrone exhibits pro-inflammatory and proliferative responses while 2-hydroxyoestradiol and 2-methoxyoestradiol exhibit anti-inflammatory and antiproliferative responses.^13, 14^ In IPAH, there is a higher preference for estradiol to be metabolized into 16α-hydroxyoestrone, a metabolite that is converted by the CYP1B1 enzyme, which is also increased in IPAH.^3, 15^ Estrogen receptors have been linked to expression of several ECM proteins including collagens I and III.^16^ Activation of these receptors has also been associated with other ECM mediators, such as LOX, MMP2, MMP9, HIF1α, and factors linked with the mechanosensitive YAP/TAZ pathway.^17–23^

The few currently approved medications for IPAH all focus on treating vasoconstriction.^13, 24, 25^ Acknowledging the importance of estrogen signaling in disease progression, tamoxifen, a selective estrogen receptor blocker, is currently in phase two clinical trials for IPAH (ClinicalTrials.gov identifier NCT03528902). Tamoxifen binds to estrogen receptors α and β, blocking activation of these receptors. Conversely, tamoxifen can act as a G protein-coupled estrogen receptor (GPER) agonist, increasing activation of these receptors.^26, 27^ In human pulmonary artery, ERα is the predominantly expressed receptor and levels increase in IPAH, with ERβ expression also observed particularly in the endothelial layer, while GPER expression is rarer, suggesting that overall in pulmonary artery tamoxifen would reduce estrogen signaling activity.^28^ Studies have shown that modulating estrogen receptors with tamoxifen inhibited pulmonary artery smooth muscle cell (PASMC) proliferation and migration.^24^ It also had anti-inflammatory effects within the pulmonary arteries.^24^ In breast cancer and hepatocellular carcinoma models, tamoxifen was found to decrease HIF1a, LOX, and YAP pathways.^17^ Tamoxifen is being studied as an antifibrotic because it can modulate the AKT pathway, a signal transduction pathway that regulates cell growth, survival, metabolism, and motility. Tamoxifen downregulates p-AKT and its downstream protein expression, which then inhibits fibroblast proliferation, and is being studied to treat epidural fibrosis, esophageal strictures, idiopathic retroperitoneal fibrosis, and renal tubulointerstitial fibrosis.^29–32^ In fibroblasts, tamoxifen has also been found to mostly inhibit estrogen signaling, subsequently inhibiting TGFβ-mediated fibroblast activation.^33^ Endoxifen is the most active secondary metabolite of tamoxifen. Conditions such as gynecologic tumors, desmoid tumors, breast cancers, and hormone receptor-positive solid tumors that use tamoxifen as a treatment are currently studying endoxifen, which has a higher affinity for estrogen receptors, as a new therapeutic option.^34, 35^

Pulmonary vascular remodeling involves multiple cell types and is driven by dysregulated extracellular matrix (ECM) dynamics and mechanosignaling. In healthy tissue, hPAAFs regulate ECM homeostasis through the balanced synthesis and degradation of proteins, glycosaminoglycans, and other molecules. However, during IPAH disease progression, hPAAF activation becomes dysregulated, leading to abnormal proliferation, apoptosis resistance, excessive ECM production, increased stiffness, and pro-inflammatory behavior.^8, 9, 36, 37^ Vascular remodeling in IPAH is characterized by an imbalance between ECM-degrading enzymes and their regulators, resulting in increased fibrillar ECM deposition, crosslinking, and elastin degradation.^38^

This buildup of highly crosslinked ECM increases vascular stiffness, activating three major artery cell types (hPAAFs, PASMCs, and pulmonary artery endothelial cells) through mechanosignaling pathways. Tunable stiffness hydrogels have become a valuable tool for studying these mechanosignaling effects in vitro, mainly in 2D systems where cells are grown on defined stiffness substrates. Studies have shown that pulmonary artery cells grown on stiff surfaces expressed elevated levels of ECM factors and mechanosignaling molecules, including Col1a1, Col3a1, CTGF, LOX, YAP, and miR-103/301, compared to cells cultured on softer surfaces.^37, 39, 40^ hPAAFs showed increased alpha-smooth muscle actin (αSMA) expression when cultured on stiff matrices, indicating a more activated, myofibroblast-like state.^37^

Sex differences also influence hPAAF activation in response to mechanical cues and circulating hormones. Previous work from our lab investigated the phenotype of hPAAFs cultured on soft and dynamically stiffened hydrogels in medium containing human serum pooled by age and sex.^11^ Female hPAAFs exhibited greater activation on both soft and stiff matrices, except when cultured in serum with higher concentrations of estradiol, typical of females under age 50. Male hPAAFs showed increased activation, specifically on stiffened substrates, regardless of circulating sex hormone levels.^11^ These findings emphasize that both microenvironmental stiffness and circulating sex hormones must be considered to model sex differences in hPAAF activation in vitro accurately.

The current study builds on this work by incorporating control or IPAH donor female hPAAFs into 3D-bioprinted hydrogel constructs that mimic the adventitial layer of the pulmonary artery and culturing these models in sex-specific human serum. Bioprinted constructs were designed to be approximately 4-mm in diameter with 300-500 *μ*m wall thickness replicating the dimensions of third to fourth generation pulmonary arteries, which are commonly affected in IPAH.^10, 41^ Constructs were bioprinted using hydrogel bioinks comprised of a hydrolytically stable, alpha-methacrylated poly(ethylene) glycol (PEGαMA) crosslinked with a cell-degradable peptide sequence and decorated with a cell-adhesive peptide (CGRGDS) to allow hPAAFs to attach to and dynamically remodel the microenvironment. Hydrogels were designed to initially mimic the microenvironmental stiffness of healthy pulmonary arterial tissue (E = 1–5 kPa) and then be dynamically stiffened to within the range of hypertensive pulmonary arterial tissues (E > 10 kPa).^7^ Through the duration of the experiment, samples were cultured in female human serum collected from donors under 50 years old to provide physiologically relevant biochemical cues, including circulating estradiol. Stiffened constructs showed increased activation levels, with IPAH donor fibroblasts showing enhanced activation by protein and gene expression. The addition of endoxifen to hypertensive models reduced expression of fibroblast activation markers. Collectively, these results present an *in vitro* method for studying the crosstalk between cell origin, estrogen signaling, and matrix stiffness in the adventitial layer of the pulmonary artery and that sex-specific *in vitro* models can be useful tools for discovering and validating drugs for populations that are more susceptible to IPAH.

## 2. Materials and Methods

### 2.1 PEGαMA Synthesis and Hydrogel Characterization

PEGαMA was synthesized as previously described.^42, 43^ To summarize, poly(ethylene glycol)-hydroxyl (PEG-OH; 8-arm, 10 kg mol-1; JenKem Technology) was dissolved in tetrahydrofuran (THF; Sigma-Aldrich) and a 3 molar excess to hydroxyls of sodium hydride was added to the reaction to deprotonate hydroxyl groups. Then, a 6 molar excess to hydroxyl groups of ethyl 2-(bromomethyl)acrylate (EBrMA; Ambeed, inc.) was added dropwise to the reaction and was stirred for 48 h protected from light.

Acetic acid was used to neutralize the reaction, and the solution was filtered through Celite 545 (EMD Millipore) and concentrated by rotary evaporation twice. Ice-cold diethyl ether was added to the concentrated solution and allowed to sit overnight at 4°C. The precipitated product was dialyzed against deionized water for four days, then flash-frozen and lyophilized until dry. A small amount of product was dissolved in chloroform-d (CDCl3) for proton nuclear magnetic resonance (1H NMR). 284 scans were acquired on a 300 MHz Bruker DPX-400 FT NMR spectrometer with a 2.5-second relaxation delay as previously described.^42, 43^ 1H NMR (300 MHz, CDCl3): d (ppm) 1.36 (t, 3H, CH3–), 3.71 (s, 114H, PEG CH2-CH2), 4.29 (t, s, 4H, –CH2–C(O)–O–O, –O–CH2–C(=CH2)–), 5.93 (q, 1H, –C = CH2), 6.34 (q, 1H,–C = CH2).^44^ PEGαMA with functionalization over 90%, as measured by comparing the αMA vinyl end group peak to that of the PEG backbone, was used in subsequent experiments (Figure S1).

Hydrogel precursor solutions were prepared by combining 17.2 weight % (wt%) PEGαMA with dithiothreitol (DTT; Sigma-Aldrich) and the MMP-degradable peptide crosslinker KCGPQGIWGQGCK (70:30 crosslinker ratio; GL Biochem) off-stoichiometry with a [thiol]:[ene] ratio of 0.375. This peptide sequence can be degraded by MMP1, MMP2, MMP3, MMP8, and MMP9,^45^ however, it is most responsive to MMP2. Hydrogels contained 2 mM of the adhesive peptide CGRGDS (GL Biochem), which mimics cellular binding cites on the protein fibronectin, and 2.5 wt% poly(ethylene oxide) (PEO; 400 kg mol-1; Sigma-Aldrich) to enhance shear-thinning properties of the bioink and facilitate 3D bioprinting.^10^

Hydrogels were polymerized through a base-catalyzed Michael addition reaction to make initially soft hydrogels. The hydrogels for rheological evaluation were formed by placing 40 uL drops of hydrogel precursor solution between two parafilm-covered glass slides with a 1-mm silicone gasket. After polymerization, 1-mm thick hydrogels that were approximately 8 mm in diameter were formed. The hydrogels were allowed to equilibrate overnight in PBS with 2.2 mM lithium phenyl-2,4,6-trimethylbenzoylphosphinate (LAP; Sigma-Aldrich) photoinitiator. The hydrogels intended to be stiffened were exposed to 365 nm light, 10 mW cm-2 (Omnicure, Lumen Dynamics) for 5 minutes initiating a homopolymerization reaction between the unreacted α-methacrylate groups, dynamically increasing the average elastic modulus of the hydrogels.

Rheology was performed as previously described.^10, 11^ Storage modulus (G’) was measured on a Discovery HR-2 rheometer (TA Instruments) using an 8-mm parallel plate geometry at 37°C. After the geometry was lowered to an axial force of 0.03 N, the gap distance was measured and was adjusted starting at 25% compression until the storage modulus (G’) stabilized under compression. The percent compression at which G’ plateaued was used for measurement samples. An oscillatory frequency sweep with oscillatory shear at 1% strain through a 1-100 rad/s angular frequency range was used to measure G’. Using rubber elastic theory, the elastic modulus of elastic hydrogel polymer networks was calculated using a Poisson’s ratio of 0.5.^46^

### 2.2 Cell Culture

hPAAFs were acquired from the Pulmonary Hypertension Breakthrough Initiative (PHBI). Age-matched cells from IPAH patients and control donors (Table 1) were both cultured according to supplier instructions. Briefly, cells were expanded at 37°C and 5% CO2 on tissue-culture-treated polystyrene until 90-95% confluency and then passaged for 3D bioprinting. Growth medium consisted of SmGM basal medium (Lonza CC-3181) with 5% fetal bovine serum (FBS), 0.1% insulin, 0.2% human fibroblast growth factor, 0.1% gentamicin sulfate - amphotericin, and 0.1% human epidermal growth factor (Lonza CC-4149).

**Table 1.** PHBI hPAAF donor information.

| PHBI App # | Disease | Age |
| --- | --- | --- |
| PHBI-AH-014 | Normal | 28 |
| PHBI-BA-054 | Normal | 33 |
| PHBI-UA-015 | Normal | 36 |
| PHBI-BA-019 | IPAH | 29 |
| PHBI-VA-011 | IPAH | 32 |
| PHBI-BA-004 | IPAH | 39 |

### 2.3 3D Bioprinting

To model third to fourth generation human pulmonary arteries, a 3D computer-aided design (CAD) model with 4-mm height, 4-mm inner diameter, and 300-µm thickness was developed in Fusion 360.^41, 47^ Then, the design model was sliced into G-code using Slic3r software and uploaded to Pronterface, the software interface for the modified Mini Lulzbot 2 (Aleph Objects, Inc.) 3D printer that was used throughout these studies.^48^ All 3D-bioprinted constructs used this model and were printed using a slightly adapted Freeform Reversible Embedding of Suspended Hydrogels (FRESH) 3D bioprinting method.^10, 49^

Briefly, 1 g of LifeSupport gelatin microparticle slurry (Advanced BioMatrix, cat. #5244-8GM) was rehydrated with 40 mL of cold serum-free, phenol-free basal medium (Lonza), pH adjusted to 10.5 and then compacted within a 50 mL conical tube.

Compacting was completed by centrifuging the solution for 5 min at 2000 x g, pouring off the supernatant, gently tapping the conical tube 15 times, inverting the conical tube back and forth for 10 s, centrifuging for another 5 min at 2000 x g, pouring off any remaining supernatant, and then storing at 4 °C for approximately 1 h before use.

In parallel, stock solutions of hydrogel precursors were prepared. 250 mM DTT, 100 mM MMP-degradable peptide crosslinker, and 250 mM CGRGDS were individually resuspended in 20 mM TCEP (pH 7), each sonicated at 40 °C for 5 min, and then were vortexed. The 17.2 wt% PEGαMA was resuspended in sterile complete cell culture medium (pH 7) at a concentration of 23.13 mM and then briefly vortexed. A 15 wt% PEO in PBS solution was prepared in advance. Once all components were completely in solution, the correct proportions of each were combined with hPAAFs (passages 3-6) to produce the final bioink (pH ∼6.5) formulation (17.2 wt% PEGαMA, 6.45 mM DTT, 15 mM MMP-degradable peptide crosslinker, 2 mM CGRGDS, and 2.5 wt% PEO). The bioink was then transferred into the printing syringe and the syringe was secured and oriented within the bioprinter.

Just prior to printing, the tip of the conical tube containing the support bath material was manually cut off with a razor and the support bath material was transferred into the wells of 48-well tissue culture plastic plates using a syringe plunger.^48^ Bioprinted constructs were produced and left to polymerize for 1 h at room temperature in a biosafety cabinet.^48^ Next, the plates containing the constructs were placed in a tissue culture incubator (37 °C and 5% CO2) and left overnight. The next day, melted support slurry was aspirated from each well and replaced with activation medium: phenol-free basal medium containing 1% female human serum (Innovative Research) and 75 µg mL-1 L-ascorbic acid (Sigma-Aldrich) to promote mature collagen matrix maintenance by fibroblasts ^50^. Phenol-free medium was used to eliminate the estrogenic effects of phenol red.^51, 52^ Female human serum was pooled from three donors under the age of 50 (Table 2) to reduce biological variability.

**Table 2.** Female human serum information.^11^.

| Age | Estradiol |  | Progesterone |  | Testosterone |  |
| --- | --- | --- | --- | --- | --- | --- |
|  | (pg/mL) | Average | (ng/mL) | Average | (ng/mL) | Average |
| 43 | 64 | 115.7 | 0.1 | 0.17 | 0.339 | 0.353 |
| 36 | 54 |  | 0.1 |  | 0.402 |  |
| 29 | 229 |  | 0.3 |  | 0.319 |  |

### 2.4 3D Model Culture and Stiffening

Following 3D-bioprinting activation medium (phenol-free basal medium containing 1% female human serum (Innovative Research) and 75 µg mL-1 L-ascorbic acid (Sigma-Aldrich)) was replenished on day three. To facilitate stiffening of hydrogel constructs, 2.2 mM LAP was added to the experimental medium for all samples on day six. On day seven, half of the constructs were stiffened by exposure to 10 mW cm-2 365-nm light for 5 min. All samples were processed for protein and gene expression analyses on day nine.

### 2.5 Evaluation of Cellular Responses to a Selective Estrogen Receptor Modulator

For studies involving endoxifen treatment, either 1 uM endoxifen or vehicle (EtOH:PBS 1:2) was supplemented into activation medium starting on day 1. As an active metabolite of tamoxifen, endoxifen supplementation should robustly and selectively mediate estrogen receptors in hPAAFs.

### 2.5 Immunofluorescent Staining

Samples were fixed, embedded, cryosectioned, and immunostained as previously described ^10^. To summarize, constructs were fixed in 4% paraformaldehyde (PFA; Fisher Scientific) for 30 minutes, quenched in 100 mM glycine in PBS for an additional 30 minutes, and then soaked in optimal cutting temperature compound (OCT; Fisher Scientific) overnight in a hydrated chamber. OCT-infused constructs were then flash-frozen in a liquid-nitrogen-chilled isopentane bath and stored at -80°C. Constructs were cryosectioned on a Leica CM1850 cryostat at -22°C into 10 µm-thick sections and attached to positively charged glass microscope slides. Cryosections were fixed with ice-cold acetone for 15 minutes and then rinsed with water to remove OCT. After drying, sections were outlined with a hydrophobic pen and blocked with 5% bovine serum albumin (BSA; Fisher Scientific) in PBS for one hour at room temperature. Primary antibodies (Table S1) were diluted in 3% BSA/PBS and incubated with samples overnight before washing 3x in 0.1% Tween 20 (Sigma-Aldrich) in PBS. Samples were incubated with secondary antibodies in 3% BSA/PBS (Table S1) for one hour, protected from light and again washed 3x in 0.1% Tween 20/PBS. Samples were counterstained in 10 µg/ml Hoechst nuclear dye (Tocris Bioscience) in PBS for 15 minutes. Slices were imaged on an Olympus BX63 epifluorescent microscope by imaging 3-5 random fields of view per section.

### 2.6 Quantitative Real-Time Polymerase Chain Reaction (qRT-PCR)

Constructs were transferred to sterile microcentrifuge tubes and flash-frozen in liquid nitrogen. TRIzol (Fisher Scientific) with 50 ug/mL tRNA (from *s. cerevisiae*; Sigma-Aldrich) was added to frozen constructs. Samples were dissociated with an RNase-free plastic pestle, pipetted to homogenize, and incubated 5 minutes at room temperature. The organic phase, interphase, and aqueous phase were separated by adding 100 µL 1-bromo-3-chloropropane (BCP; Fisher Scientific) and incubating for 15 minutes at room temperature. Samples were cooled on ice for 1 minute and centrifuged at 12,000 × g for 15 minutes. The top aqueous RNA phase was transferred to a fresh RNase-free microcentrifuge tube and mixed 50:50 with 100% ethanol. Samples were run through RNeasy Mini columns (Qiagen) for purification according to the manufacturer’s instructions. cDNA was synthesized using the High-Capacity cDNA Reverse Transcription Kit (Applied Biosystems) using the manufacturer’s protocol. qRT-PCR was run using iTaq Universal SYBR Green Supermix (Bio-Rad) following manufacturer protocols on a Bio-Rad CFX96 thermal cycler (Bio-Rad). Human primers were acquired from Integrated DNA Technologies (**Error! Reference source not found.**S2). Data were normalized to the reference gene ribosomal protein L30 (RPL30). Data points with a Ct less than 35 were considered for ΔCt quantification analysis.

### 2.7 Statistical Methods

Sample means from three or more groups were compared using ordinary two-way ANOVA with α = 0.05 and Tukey tests for multiple comparisons. F-tests were conducted to determine equal sample variance between donor results, and donor data were pooled when these sample variances did not differ. Outliers were removed using the ROUT method with Q = 5%. All statistical tests were conducted in GraphPad Prism (GraphPad Software). Unless otherwise reported, data are presented as mean ± standard error of the mean (SEM) for three biological replicates of each condition.

## 3. Results and Discussion

### 3.1 3D Bioprinted Hydrogel Synthesis and Characterization

Distal pulmonary artery stiffening is an early regulator of vascular remodeling in IPAH.^7^ As the disease progresses, the thickness of the adventitial layer thickness of these smaller pulmonary arteries has been shown to increase 28% and is primarily attributed due to increased collagen deposition.^53^ Therefore, to accurately produce physiologically relevant models of diseased pulmonary artery adventitia, tubular constructs with 4-mm diameter and 300-µm thickness walls containing female control or IPAH fibroblasts were 3D bioprinted.^41, 47^ Freeform reversible embedding of suspended hydrogels (FRESH) is a 3D bioprinting technique developed by the Feinberg lab that entails depositing a hydrogel precursor solution in a thermoreversible support bath. This technique allows for the formation of complex 3D structures that could not be achieved by conventional 3D bioprinting and was used here to create adventitial models.^49^

Most *in vitro* pulmonary artery models investigate how vascular shape or stiffness effect cellular response, but few models specifically replicate the adventitial layer of pulmonary arteries. A study by Jin et al. developed a self-folding cell-laden model that formed tubular constructs. The size of the tube was controlled to make models of both distal and proximal pulmonary arteries. The backbone of the model was a thin film of silicone oxide and silicone dioxide. Smooth muscle cells and endothelial cells were seeded on the thin film prior to the films folding, creating a tubular model that consisted of both smooth muscle cells and endothelial cells, but this model was not at physiologic stiffness with a stiffness of 1-200 GPa and did not incorporate the adventitial layer of the pulmonary arteries.^54^ Another tubular model used a double network hydrogel, where gelatin-methacrylate and sodium alginate were used to print a double layered hydrogel tube that incorporated smooth muscle cells and endothelial cells. This allowed for the formation of a functional artery or vain model that was of physiologically relevant stiffness, but did not incorporate the adventitial layer on the artery models.^55^ In comparison, the few published *in vitro* models that do study the adventitial layer of the pulmonary artery grew these cells on physiologically relevant 2D stiffness biomaterials,^11, 37^ but lacked 3D geometries.

To overcome this limitation, we leveraged the FRESH bioprinting technique and produced an adventitia model that incorporates both physiologically relevant stiffness and shape. This involved making a bioink consisting of a PEGαMA backbone with two crosslinkers (DTT and MMP-degradable peptide crosslinker), a cell adhesion peptide (CGRGDS), and the cells of interest. PEO was incorporated to allow for shear thinning properties of the bioink. The bioink would be bioprinted into a gelatin microparticle bath which allows for the printing of complex hydrogel structures. The bioink is kept acidic (pH 6.5 - 7) and printed into a basic gelatin bath (pH 10.5) to polymerize hydrogels through a base-catalyzed Michael Addition reaction. These hydrogels were polymerized off-stoichiometry (0.375 thiols to αMA moieties) to form soft hydrogels. Addition of LAP photoinitator and UV light (365 nm) initiated a homopolymerization reaction between the unreacted αMA moieties (Figure 1A). To mimic the stiffnesses observed in healthy and hypertensive pulmonary artery tissue, hydrogels were designed to start within the stiffness range of healthy pulmonary arteries (1-5 kPa) and stiffen into the range of diseased pulmonary arteries (< 10 kPa)^7^. Soft hydrogels had an elastic modulus of 4.47 ± 0.25 kPa and stiffened hydrogels had an elastic modulus of 19.00 ± 2.93 kPa (Figure 1B).

**Figure 1.**
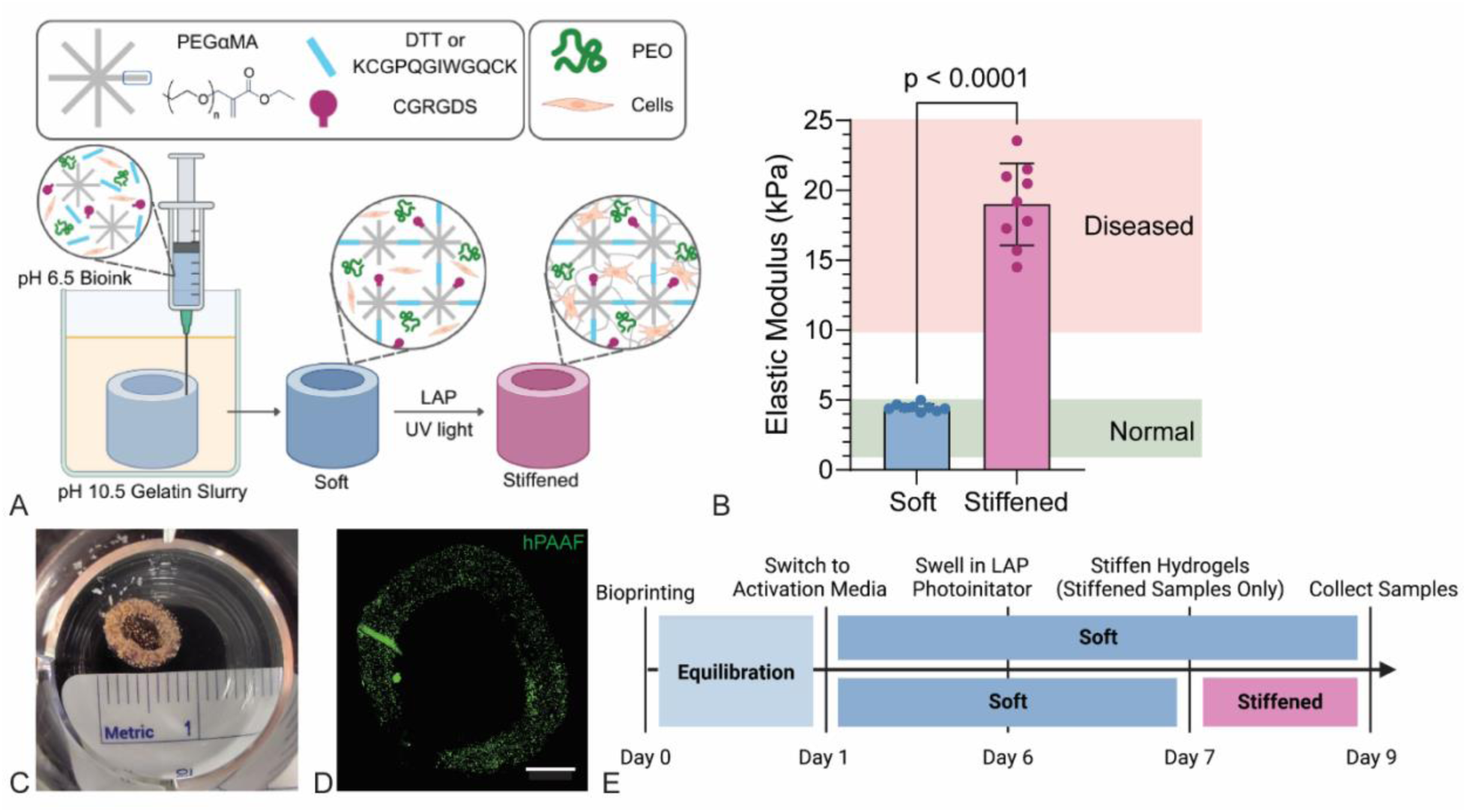
3D-bioprinted PEGαMA hydrogels were designed to dynamically stiffen mimicking changes in IPAH tissue. A) Schematic of the hydrogel formulation containing hydrogel components (PEGαMA, DTT, MMP degradable crosslinker, and CGRGDS) and components in suspension within hydrogels (PEO and cells). Hydrogel components were maintained at pH 6.5 and 3D-bioprinted into a basic (pH 10.5) gelatin slurry bath initiating a base-catalyzed Michael Addition reaction to form soft hydrogels constructs.

hPAAFs from control donors or donors with IPAH were bioprinted into soft hydrogels and later stiffened to investigate the influence of cell origin on fibroblast response and how microenvironmental stiffness effects the response. Bioprinted constructs were equilibrated overnight in the gelatin slurry containing growth medium, after which the tubes showed an outer diameter of approximately 7 mm and a thickness of 1 mm (Fig 1C), with hPAAFs evenly distributed throughout (Fig 1D). The next day, medium was changed to activation medium containing 1% female human serum pooled from three donors less than 50 years old (Table 2).^11^ On day six, LAP photoinitator was added to all samples and on day seven, half of the samples where stiffened with 365 nm light. Samples were grown for two more days (day nine) before samples were collected for analysis (Figure 1E). To explore the impact of estrogen signaling on hPAAF activation response, endoxifen, a selective estrogen receptor modulator, or a vehicle was applied to samples starting on day one.

Incubation in LAP and exposure to UV light initiated a homopolymerization reaction to stiffen the hydrogel constructs. B) Rheological measurements showed the average elastic modulus (E) of soft hydrogels (E = 4.47 ± 0.25 kPa) was within the range of healthy pulmonary arteries (1–5 kPa) and the average elastic modulus of stiffened hydrogels (E = 19.00 ± 2.93 kPa) increased into the range of pathologic pulmonary arteries (>10 kPa). Columns represent mean ± SD, n = 9. C) Color image of a bioprinted tube in a 48-well plate. Ruler for scale. D) Fluorescent image of hPAAFs labeled with CellTracker Green (ThermoFisher) inside a bioprinted construct. Scale = 2 mm. E) Schematic representation of the nine-day timeline used for cell experiments with stiffening on day seven.

### 3.2 Fibroblast activation phenotype is controlled by both extrinsic and intrinsic factors

Most *in vitro* model systems rely on fetal bovine serum (FBS) as a nutrient source for cells. However, FBS may contain a unique blend of hormones, that varies by lot and is undisclosed by most manufacturers.^56^ Most notable the samples tested by our lab previously contained extremely high progesterone levels, likely from a maternal source.^11^ Cells exposed to FBS *in vitro* may often be exposed to non-physiologically hormone signaling. To address this limitation, we cultured hPAAF-containing bioprints in medium supplemented with age- and sex-matched human serum. We specifically aimed to build a model of hPAAF activation in young (age < 50) female samples to match the demographic most likely to be diagnosed with IPAH. Human serum was pooled from three female donors ages 29-43, and contained physiologic levels of estradiol, progesterone, and testosterone (Table 2). Female patient-derived fibroblasts were sourced from control patients or IPAH arteries. 3D bioprinted samples either remained soft to mimic health pulmonary arteries or were dynamically stiffened to investigate the relative contributions of cell source and microenvironmental stiffness to changes in cellular phenotype. Activation was assessed by both immunostaining and qPCR for a panel of IPAH-related factors.

First, staining was performed for αSMA, a classical marker for activated myofibroblasts that are observed at sites of active tissue remodeling in many injury responses and disease states, including IPAH. Next, staining for collagens I and III quantified accumulation and increased crosslinking of fibrillar collagen, a hallmark of IPAH in both proximal and distal arteries.^57^ In cardiac fibroblasts, collagen expression has been shown to be responsive to estradiol signaling in a sex specific manner, and the ratios of collagen I to collagen III changed with fibrosis, so both proteins were examined in our female-specific model. Stiffness significantly increased αSMA expression in both control and IPAH cells, with approximately 60% αSMA-positive hPAAFs in soft conditions and nearly 80% in stiffened (Figure 2A-B). Conversely, collagen expression was largely controlled by cell source, with around 30-40% of IPAH cells expressing either intracellular collagen I or collagen III, compared to only 10% of control cells (Figure 2C-D). Expression trends of collagens I and III were similar, without significant changes in the ratio between the two. Lung fibroblast heterogeneity has been well established in many chronic lung diseases, and gene expression profiling suggests that αSMA and collagen I expression can serve as markers for unique fibroblast subsets, with collagen-positive fibroblasts playing roles in ECM synthesis, while contractile αSMA-positive fibroblasts are more prominent in ECM remodeling.^58, 59^ Less research has been performed on hPAAF heterogeneity specifically, but one study demonstrated that unique hPAAF subsets can be identified by morphology, marker expression, and responses to stimuli such as hypoxia or growth factors, and that these subtypes are likely to play unique functional roles in both healthy and diseased artery.^60^ Our data suggest that hPAAFs may separate into similar functional subtypes as other pulmonary fibroblast groupings, and that both cell intrinsic factors (cell source) and cell extrinsic factors (microenvironmental stiffness), contribute to different fibroblast activation pathways, with stiffness strongly disposing towards a contractile αSMA-positive phenotype, while cellular disease state promotes a collagen-expressing ECM synthetic phenotype.

**Figure 2.**
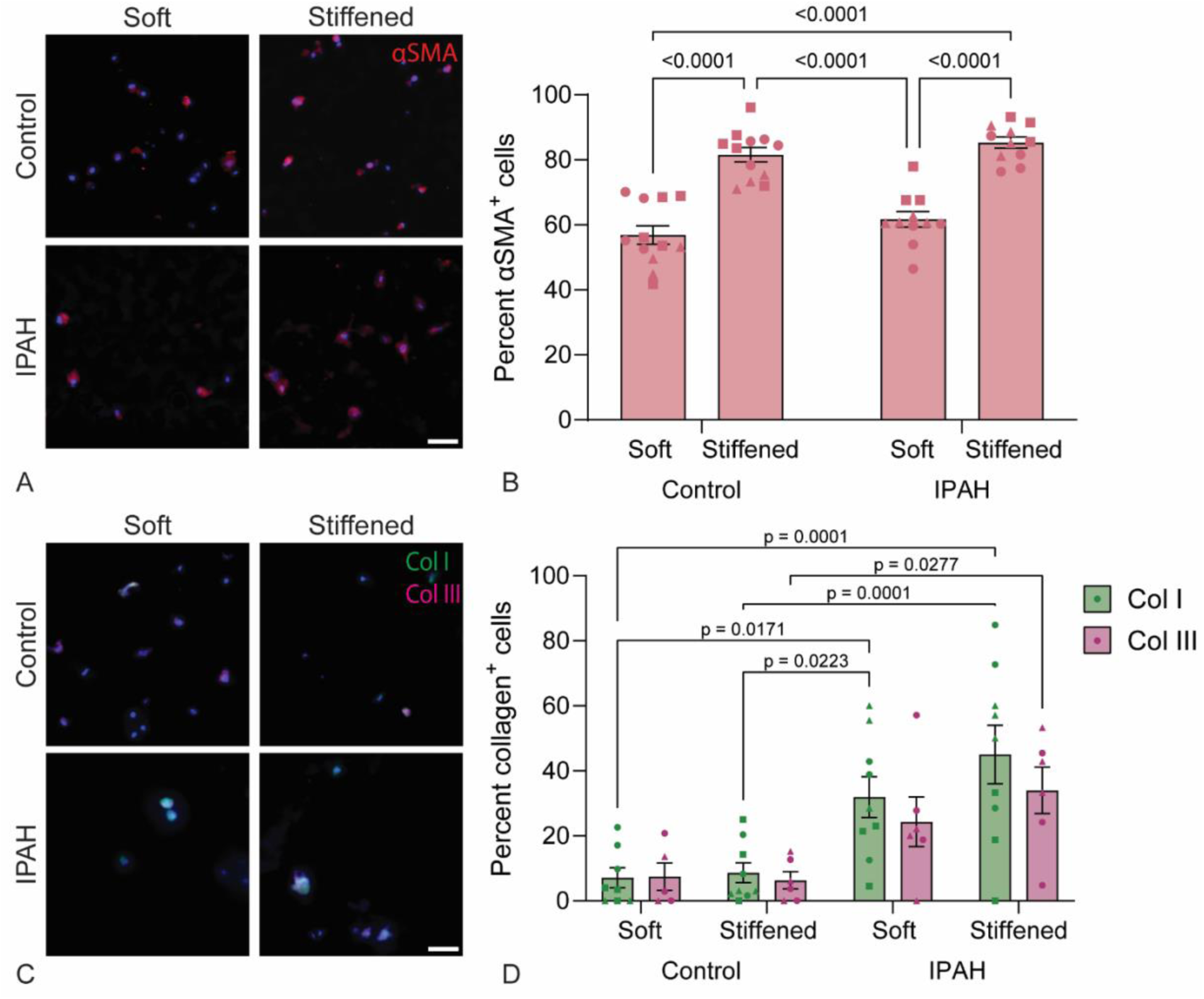
Female hPAAF activation was modulated by microenvironmental stiffness and cellular disease state. A) Activation of control and IPAH patient-derived female hPAAFs in soft and stiffened constructs measured by αSMA expression. Cells in stiffened constructs were more likely to be αSMA^+^ than cells in soft constructs in both control and IPAH cells. B) Activation of control and IPAH patient-derived female hPAAFs in soft and stiffened constructs measured by collagen I and III expression. Cells from IPAH patients were more likely to express both collagens than those from control patients. Data are presented as mean ± SEM, n = 5-9 bioprints derived from 3 patient cell lines (lines identified by dot shape). P-values reported from two-way ANOVA with Tukey tests for multiple comparisons. Scale bar = 50 µm.

To assess hPAAF activation phenotype in more detail, qPCR quantified expression of a panel of IPAH-associated genes, including several related to mechanosignaling, estrogen signaling, and the links between these two pathways (Figure 3). ACTA2, CTGF, HIF1A, and TGFA are all involved in hPAAF activation responses. ACTA2 is expressed by activated myofibroblasts, associated with contractile responses to ECM remodeling, and highly upregulated in PAH.^8, 37, 38, 61^ HIF1A is a transcription factor that is classically activated by hypoxia, linked to estrogen signaling, and promotes cellular proliferation and apoptosis resistance.^17, 22, 23, 62, 63^ CTGF is a growth factor, expression of which has also been linked to estrogen signaling as well as the YAP/TAZ mechanosensitive pathway and is highly upregulated in PAH.^17, 36–38^ Likewise, TGFA is expressed by activated hPAAFs during PAH progression.^8, 64, 65^ ECM remodeling genes analyzed include various ECM structural components, matrix remodeling enzymes, and as CTHRC1, an extracellular protein which co-localizes with αSMA and collagen at sites of arterial injury and remodeling.^61, 66^ We also investigated estrogen signaling directly by quantifying gene expression of CYP1B1, which is the critical enzyme for estrogen synthesis, as well as the estrogen receptors ESR1, ESR2, and GPER1. ESR1 and ESR2 show sex-specific regulation in the context of PAH, with females showing increased ESR1 expression while males tend towards increased ESR2.^28^

**Figure 3.**
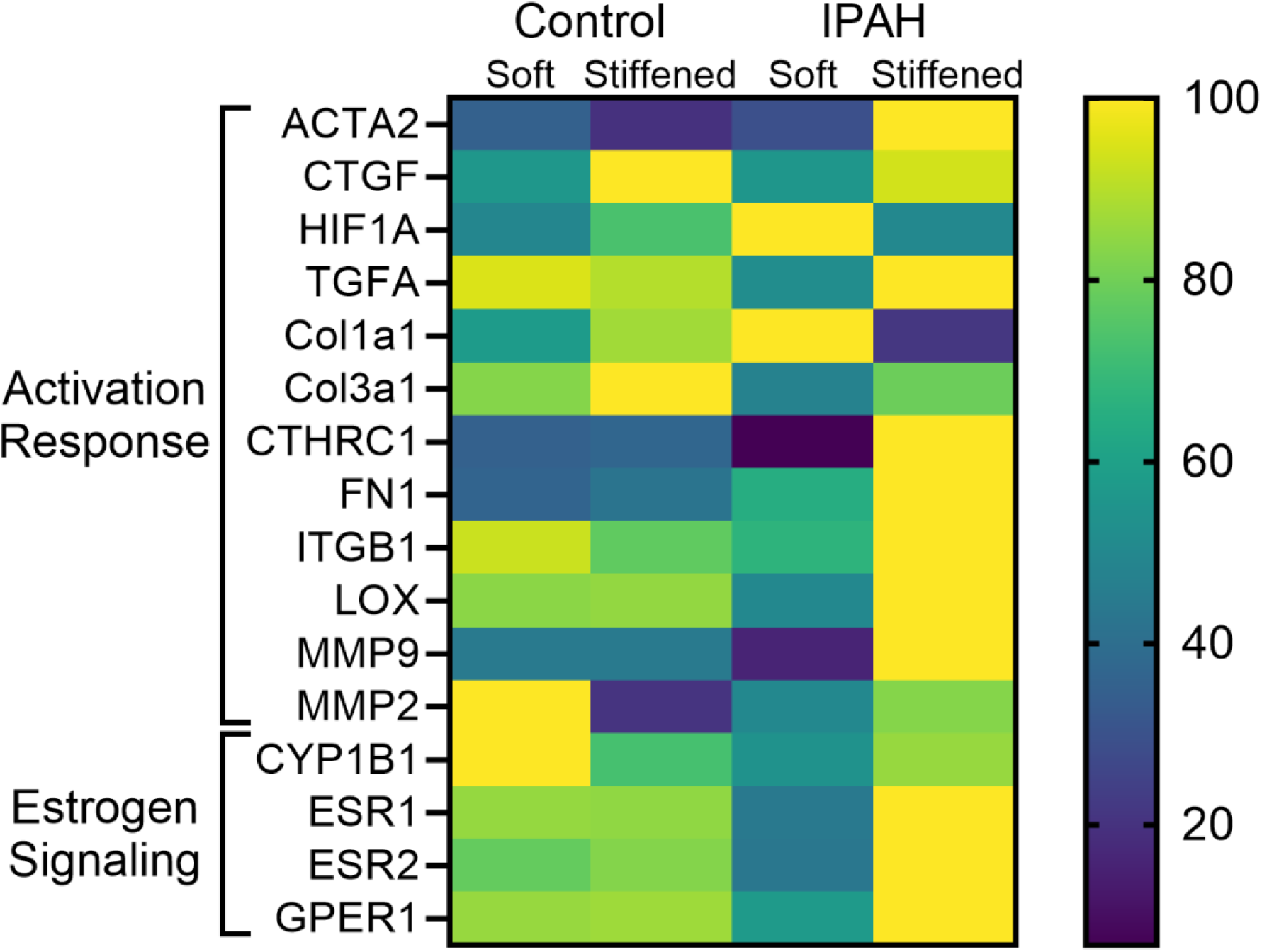
Relative expression of genes that play a role in fibroblast activation, extracellular matrix remodeling, and estrogen signaling in fibroblasts from control or IPAH donors within soft or stiffened hydrogel constructs. N = 3 technical replicates for each of 3 donor cell lines.

In sex-specific 3D bioprints, IPAH cells cultured in a stiffened microenvironment showed the highest expression disease-related genes, suggesting that disease source and stiffness can combinatorically increase fibroblast activation (Figure 3). These results were also consistent with known links between many of these factors.

Specifically, ERα activation has been reported to stimulate production of both MMP2 and MMP9, as well as FN1 and ITGB1.^18, 67–69^ HIF1α can promote ERβ expression, which then provides negative feedback to in turn reduce HIF1α.^70^ GPER signaling is upstream of CTGF, HIF1a, and the YAP/TAZ pathway, which can in turn promote expression of MMPs.^17, 23^ Our models thus replicate the links between estrogen signaling and mechanosignaling observed in IPAH. Gene expression analysis did show some unexpected results. In slight contrast to the protein-level analysis, gene-level expression of Col1a1 and Col3a1 was not highest in the IPAH stiffened constructs. This outcome could suggest changes in post-transcriptional regulation of collagen synthesis, which is supported by the high level of LOX observed in IPAH stiffened samples. Also, surprisingly, CYP1B1 expression was the highest in our control cell, soft constructs.

This result suggests that estrogen signaling is being modulated in these models more by receptor expression and activity than by estradiol synthesis.

### 3.3 Estrogen signaling mediates fibroblast activation in disease models

Next, the role of estrogen signaling in hPAAF activation was investigated by treating the most disease relevant condition (stiffened samples containing hPAAFs from IPAH female patients) with endoxifen to modulate estrogen signaling. IPAH cells were treated with either endoxifen or vehicle control for the course of a nine-day experiment, with hydrogel stiffening induced on day 7. Overall activation was assessed by co-staining for αSMA and intracellular collagen I (Fig 4A) and quantified by counting the number of cells that expressed either αSMA and/or collagen I and reporting as percent of total cells. Treatment with endoxifen overall reduced the number of activated cells in 3D lung models (Fig 4B), suggesting that estrogen signaling plays a role in fibroblast activation in PAH. Endoxifen more strongly targeted collagen-expressing fibroblasts within the activated cells, resulting in a post-treatment population with more residual αSMA-positive cells remaining. Increased expression of genes related to collagen production and ECM remodeling were also observed as a response to stiffening. These results suggest a link between ECM synthetic activation phenotypes and estrogen signaling in hPAAFs. Staining total (intra- and extracellular) collagen I was performed was performed to further probe the relationship between collagen production and estrogen signaling in 3D bioprinted models. The experimental medium contained vitamin C to allow for the deposition of a stable extracellular collagen matrix, however total collagen I staining revealed only faint signals with extracellular localization, with most of the collagen remaining in or directly around cell bodies (Fig 4D). This total collagen stain also revealed a trend towards decreased overall collagen I expression as the result of endoxifen treatment (Fig 4E).

**Figure 4.**
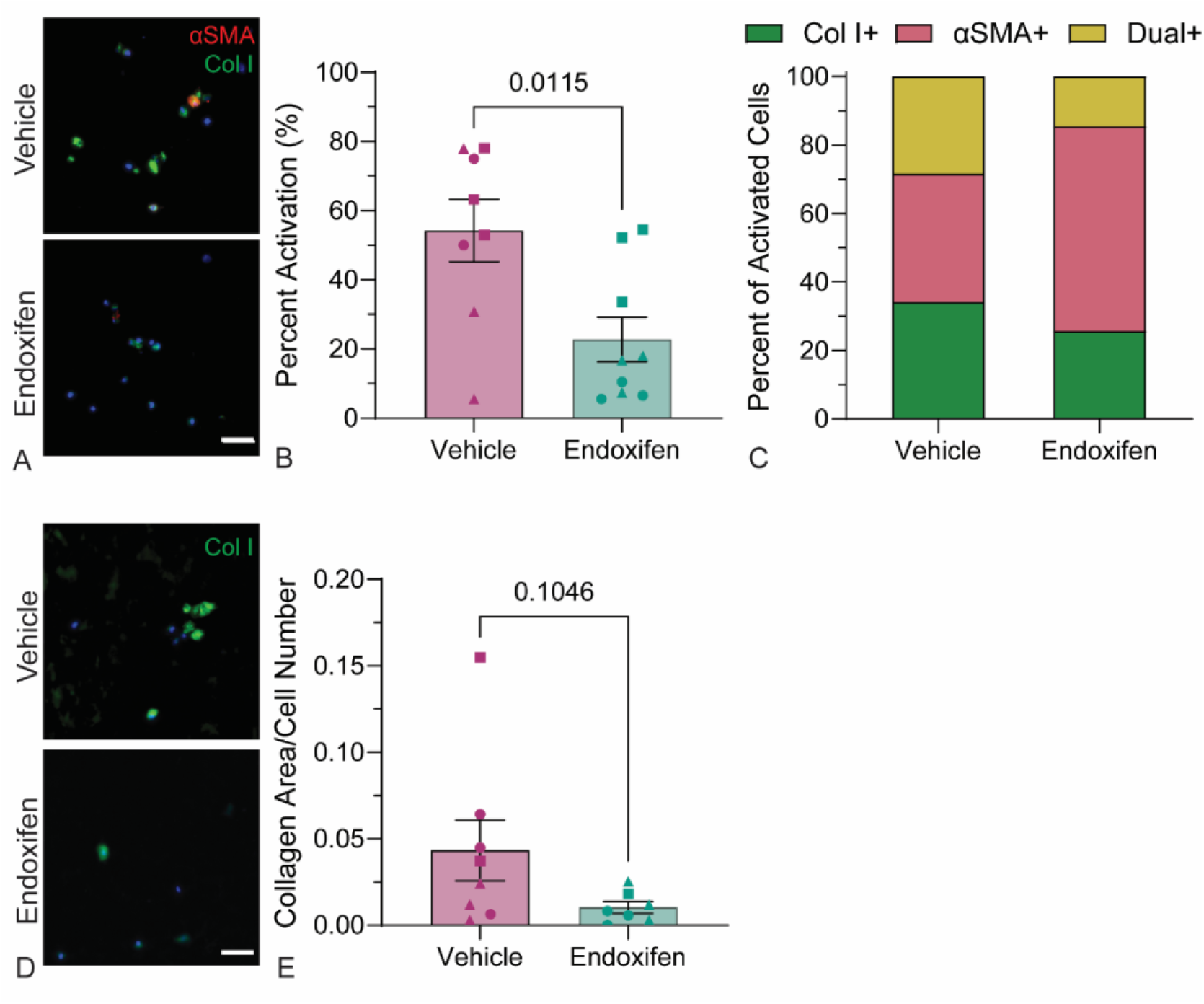
Endoxifen reduced fibroblast activation in 3D-bioprinted pulmonary artery adventitia models. A) Representative images of dual αSMA (red) and intracellular collagen I (green) immunostaining. Scale bar = 50 µm. B) The percent of cells expressing αSMA and/or collagen I was reduced by endoxifen treatment. Data are presented as mean ± SEM, n = 8-9 bioprints derived from 3 patient cell lines (lines identified by dot shape). P-values reported from student’s t-test. C) More αSMA^+^ cells remained post-endoxifen treatment within the total activated cells. D) Representative images of total (intracellular and extracellular) collagen I (green) immunostaining. Scale bar = 50 µm. E) Quantification of total collagen protein normalized to cell density with outliers removed by ROUT (Q = 1%), presented as mean ± SEM, n = 7-8 bioprints derived from 3 patient cell lines (lines identified by dot shape).

Gene expression was assessed by qPCR to further investigate the effects of endoxifen on hPAAF activation phenotypes. While many of the genes showed no change in response to endoxifen (Table S3), significant differences were observed in the expression of CYP1B1, ACTA2, CTHRC1, MMP2, and Col1a1 (Fig 5). Decreases in CYP1B1, ACTA2, CTHRC1, and MMP2 aligned with observations that endoxifen treatment reduced overall hPAAF activation. Specifically, decreases in CYP1B1 suggested reduced estradiol metabolism.^3^ MMP2 and MMP9 expression are both downstream of ESR1 signaling.^18^ Decreases in expression of these genes also demonstrated that endoxifen was actively blocking estrogen signaling pathways in 3D-bioprinted pulmonary artery adventitia models. Unexpectedly, gene expression of Col1a1 was highly upregulated. Given that reduced intra- and extracellular collagen I protein was measured in endoxifen-treated bioprints, these results suggest that modulating estrogen signaling may also cause dysregulation of collagen I synthesis and/or degradation. The accumulation of collagen I in the ECM is a tightly regulated multi-step process that includes cleavage and secretion of procollagen, crosslinking and assembly of collagen fibers, and balanced activity of collagen degradation and recycling pathways.^71–73^ Disruption at any one of these linked steps could result in less detectable collagen protein, even in the presence of high collagen transcript. In 3D-bioprinted pulmonary artery adventitia models, as endoxifen treatment causes an overall reduction in the number of collagen-expressing cells, the cells that remain might upregulate collagen transcription in a compensatory fashion, but if post-translational synthetic or degradation pathways are disrupted, the levels of extracellular collagen protein will remain low, as observed. The trends towards upregulation of TGFA, which promotes collagen upregulation during lung fibrosis^74^, LOX, which facilitates collagen crosslinking^75^, and ITGB1, which can bind to and help cells sense extracellular collagen^76^ could suggest compensatory mechanisms coming into play as hPAAFs in a stiff microenvironment attempt to maintain an activated phenotype even with endoxifen treatment.

**Figure 5.**
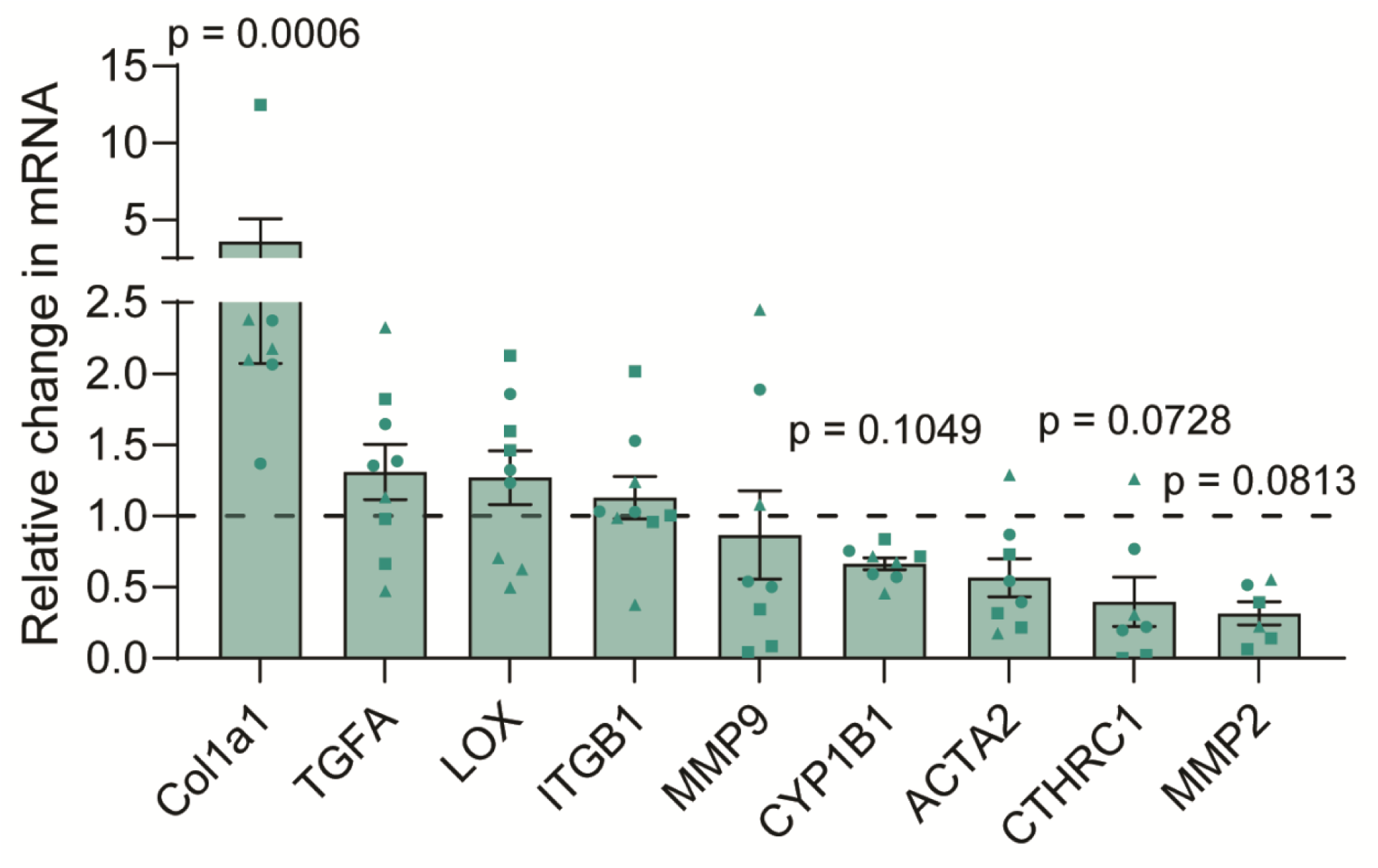
qPCR analysis of genes associated with fibroblast activation, extracellular matrix remodeling, and estrogen signaling, presented as fold change of endoxifen treated samples over vehicle treated samples. Bars represent mean ± SEM, n = 6-9 bioprints derived from 3 patient cell lines (lines identified by dot shape).

## Conclusions

*In vitro* models of PAH have traditionally relied on static, 2D culture systems and have predominately used cells of male or undisclosed sex. In this study, we shifted this paradigm by investigating the influence of microenvironmental stiffness and cellular disease state on cells that represented the patients that are most susceptible to PAH, younger females. Cell-laden, dynamically tunable hydrogels were 3D bioprinted into a geometrically relevant model of the pulmonary artery adventitial layer. These constructs were initially polymerized to replicate the modulus of healthy artery, and then dynamically stiffened to recapitulate the progressive tissue stiffening observed in IPAH. All samples were cultured using medium containing sex- and age-matched human serum. Results demonstrated that microenvironmental stiffness and tissue source (control or IPAH female donors) predisposed hPAAFs to acquire unique activated phenotypes and worked synergistically in this disease model. Estrogen signaling modulation with endoxifen, showed that estrogen signaling mediated hPAAF activation, revealing a role for hPAAF activation in the unique sex-differences clinically observed in IPAH. Collectively, these data show that estrogen-mediated mechanosignaling in hPAAFs could contribute to development of IPAH in female patients, and that 3D-bioprinted pulmonary artery adventitia models could be used to both model the estrogen paradox *in vitro* and advance drug discovery efforts for all IPAH patients. This model of pulmonary vasculature with user-controlled physiologic mechanical properties, patient-derived cells, and sex-specific culture serum will enable us to continue investigating the sequence of events that drives sex-specific responses in IPAH and drive forward precision medical treatments in this understudied population.

## Acknowledgments

hPAAFs were provided by the Pulmonary Hypertension Breakthrough Initiative (PHBI), which is funded by NHLBI #R24HL123767 and the Cardiovascular Medical Research and Education Fund (CMREF). Thanks to Claudia Staab-Weijnitz for her insights on measuring and interpreting collagen production. Schematics created with BioRender.com.

## Data Availability

The data that support the findings of this study are openly available in Mendeley Data at doi: 10.17632/grbmtp2k4w.1.

