## Supporting Information for "Female fibroblast activation is estrogen-mediated in sex-specific 3D-bioprinted pulmonary artery adventitia models"

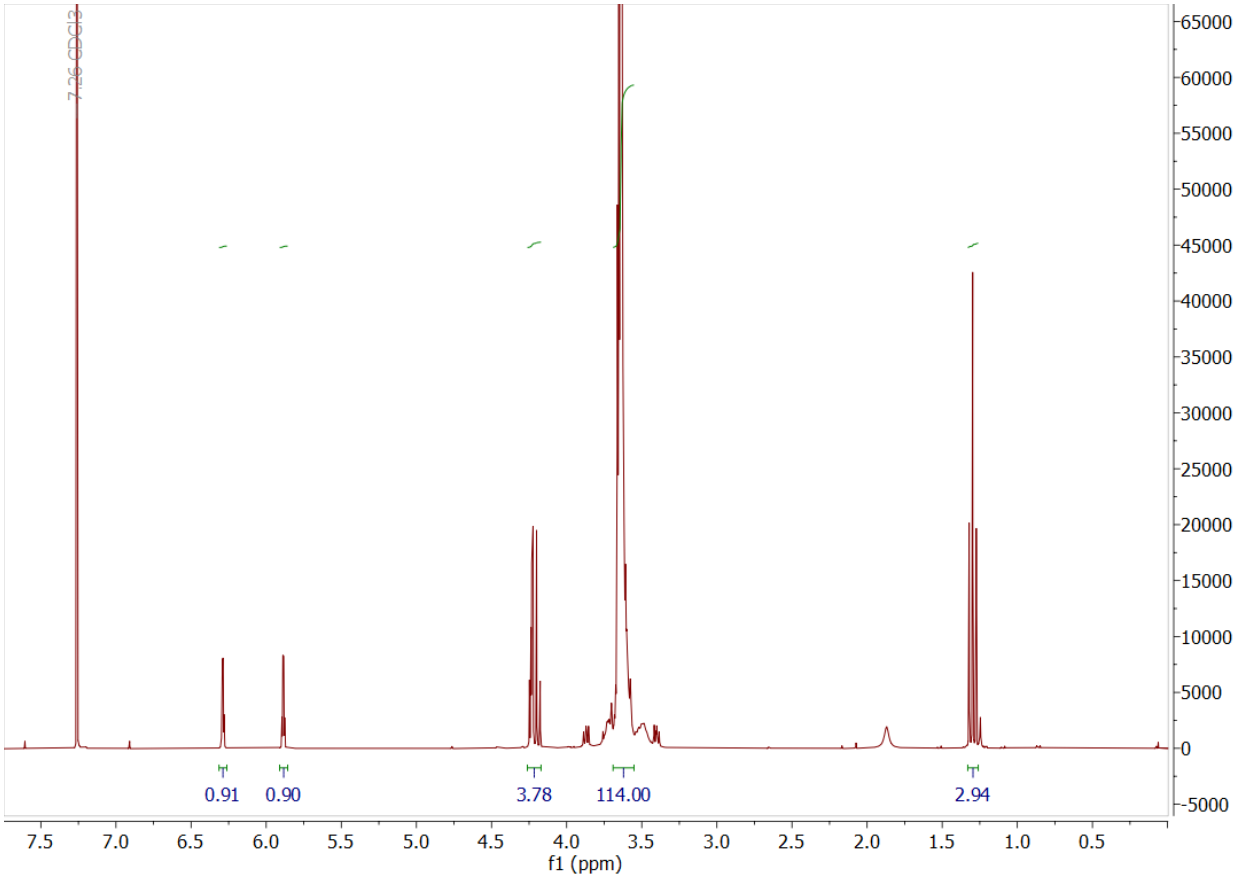


**Figure S1.** PEGαMA 1H NMR (300 MHz, CDCl3): δ (ppm) 1.36 (t, 3H, CH3–), 3.71 (s, 114H, PEG CH2-CH2), 4.29 (t, s, 4H, –CH2–C(O)–O–O, –O–CH2–C(=CH2)–), 5.93 (q, 1H, –C=CH2), 6.34 (q, 1H, –C=CH2). End group functionalization of the final PEGαMA polymer was greater than 90% by comparison of the αMA alkene end group to the PEG backbone.

**Table S1.** Antibodies used for immunofluorescent staining

| Target | Host species | Conjugate | Catalog Information |
|---|---|---|---|
| αSMA | Mouse | N/A | Fisher Scientific, MA511547 |
| Collagen I (intracellular) | Rabbit | N/A | Abcam, ab138492 |
| Collagen I (total) | Rabbit | N/A | Cell Signaling Technology, 72026S |
| Collagen III | Mouse | N/A | Sigma-Aldrich, SAB4200749 |
| GAPDH | Rabbit | N/A | Bio-Rad, AHP1628 |
| Mouse IgG | Goat | AF555 | Fisher Scientific, A21422 |
| Mouse IgG | Goat | AF488 | Fisher Scientific, A11001 |
| Rabbit IgG | Goat | AF555 | Fisher Scientific, A21428 |
| Rabbit IgG | Goat | AF488 | Fisher Scientific, A11034 |

**Table S2.** Human primers used for qRT-PCR.

| **Gene** | **Forward** | **Reverse** |
|---|---|---|
| ACTA2 | CTATGCCTCTGGACGCACAACT | CAGATCCAGACGCATGATGGCA |
| COL1A1 | TCTGCGACAACGGCAAGGTG | GACGCCGGTGGTTTCTTGGT |
| COL3A1 | ACACCGATGAGATTATGACTTCACT | TGCAGTTTCTAGCGGGGTTT |
| CTGF | AGGAGTGGGTGTGTGACGA | CCAGGCAGTTGGCTCTAATC |
| CTHRC1 | CAGACGCTGACCACGTTCC | GCATTTTAGCCGAAGTGAGCC |
| CYP1B1 | AAGAACCGCTGGGTATGGAG | TTGTGCTGCTTCTCAATTAGCG |
| ESR1 | CTGCCAAGGAGACTCGCTAC | CCTCACAGGACCAGACTCCATA |
| ESR2 | CAGGCCCACCTTCCAAGTTA | AGCCACCATGAATATCCAGCC |
| FN1 | AGGAAGCCGAGGTTTTAACTG | AGGACGCTCATAAGTGTCACC |

| | | |
|---|---|---|
| GPER1 | GGCCAATGGGACAGGTGAG | GGTGTAGAGGCACGAGAGGA |
| HIF1A | CAAAGTCGGACAGCCTCACC | GGGCTTCTTGGATGAGATTTTTCTG |
| ITGB1 | ACACCAGCAACTCTCAACATACT | TGCAGAATCCAAAGTAAATGTCCTG |
| LOX | CCATGCTATGCACCCAGTGA | GTGCCTCGGCATCCTGTAT |
| MMP2 | CAGGGAATGAGTACTGGGTCTATT | ACTCCAGTTAAAGGCAGCATCTAC |
| MMP9 | GCCACTACTGTGCCTTTGAGTC | CCCTCAGAGAATCGCCAGTACT |
| RPL30 | TCTGCTTGTACCCCAGGACG | CGAAAAAGTCGCTGGAGTCGAT |
| TGFA | GGGACCCACTGTTTCTCACAG | AGCAATTGTTCCCCTGAGCAT |

**Table S3.** Gene expression after endoxifen treatment

| Gene | Average fold change (endoxifen/vehicle) | SEM | p-value (Mann-Whitney test of endoxifen vs. vehicle) |
|---|---|---|---|
| Col1a1 | 3.563 | 1.492 | 0.0006 |
| TGFA | 1.309 | 0.1926 | 0.2973 |
| LOX | 1.270 | 0.1884 | 0.3401 |
| ITGB1 | 1.130 | 0.1494 | 0.5457 |
| Col3a1 | 1.078 | 0.1573 | 0.9314 |
| CTGF | 1.067 | 0.1557 | 0.9314 |
| GPER1 | 1.065 | 0.1466 | 0.9626 |
| ESR1 | 1.043 | 0.0259 | 0.6806 |
| FN1 | 1.039 | 0.2132 | 0.2268 |
| HIF1A | 0.995 | 0.1461 | 0.9314 |

| | | | |
|---|---|---|---|
| ESR2 | 0.9403 | 0.0886 | 0.9999 |
| MMP9 | 0.8663 | 0.3104 | 0.3213 |
| CYP1B1 | 0.6644 | 0.0425 | 0.1049 |
| ACTA2 | 0.5665 | 0.1337 | 0.3969 |
| CTHRC1 | 0.3963 | 0.1732 | 0.0728 |
| MMP2 | 0.3146 | 0.0823 | 0.0813 |
